# Movement-related modulation in mouse auditory cortex is widespread yet locally diverse

**DOI:** 10.1101/2023.07.03.547560

**Authors:** Karin Morandell, Audrey Yin, Rodrigo Triana Del Rio, David M. Schneider

## Abstract

Neurons in the mouse auditory cortex are strongly influenced by behavior, including both suppression and enhancement of sound-evoked responses during movement. The mouse auditory cortex comprises multiple fields with different roles in sound processing and distinct connectivity to movement-related centers of the brain. Here, we asked whether movement-related modulation might differ across auditory cortical fields, thereby contributing to the heterogeneity of movement-related modulation at the single-cell level. We used wide-field calcium imaging to identify distinct cortical fields followed by cellular-resolution two-photon calcium imaging to visualize the activity of layer 2/3 excitatory neurons within each field. We measured each neuron’s responses to three sound categories (pure tones, chirps, and amplitude modulated white noise) as mice rested and ran on a non-motorized treadmill. We found that individual neurons in each cortical field typically respond to just one sound category. Some neurons are only active during rest and others during locomotion, and those that are responsive across conditions retain their sound-category tuning. The effects of locomotion on sound-evoked responses vary at the single cell level, with both suppression and enhancement of neural responses, and the net modulatory effect of locomotion is largely conserved across cortical fields. Movement-related modulation in auditory cortex also reflects more complex behavioral patterns, including instantaneous running speed and non-locomotor movements such as grooming and postural adjustments, with similar patterns seen across all auditory cortical fields. Our findings underscore the complexity of movement-related modulation throughout the mouse auditory cortex and indicate that movement related modulation is a widespread phenomenon.

**SIGNIFICANCE STATEMENT:** Throughout the sensory cortex, neural activity is influenced by behavior. It remains unknown whether primary and higher-order sensory cortical centers are similarly or differentially influenced by movement. We show that movement-related modulation in the mouse auditory cortex is locally complex and heterogeneous, but that at a more macroscopic level, the net effect of movement on primary and higher-order auditory cortex is largely conserved. These data highlight the widespread nature of movement-related modulation and suggest that movement signals may inform neural computations throughout multiple nodes of the sensory cortex.

## INTRODUCTION

Neural activity in sensory cortices is influenced by movement (Schneider, Nelson, and Mooney 2014; Niell and Stryker 2010; Stringer et al. 2019; Schneider 2020). In the mouse auditory cortex, movement-related modulation manifests as changes in the spontaneous and sound-evoked responses of neurons during behaviors including locomotion, forelimb movements, and vocalizing (Schneider, Nelson, and Mooney 2014; Audette et al. 2022; Zhou et al. 2014; Rummell, Klee, and Sigurdsson 2016; Henschke, Price, and Pakan 2021; Vivaldo et al. 2022; Bigelow et al. 2019). Movement-related modulation of individual neurons is diverse within and across experimental paradigms, often leading to the generic suppression of spontaneous and sound-evoked activity (Schneider, Nelson, and Mooney 2014; Zhou et al. 2014; Bigelow et al. 2019), but also enhancement of responses (Henschke, Price, and Pakan 2021; Vivaldo et al. 2022) and acoustically selective modulation that arises with motor-sensory experience (Schneider, Sundararajan, and Mooney 2018; Rummell, Klee, and Sigurdsson 2016; Audette et al. 2022).

This diversity of movement-related modulation may reflect broad heterogeneity of neural responses throughout the auditory cortex. Alternatively, it could arise from differences in where within the auditory cortex neurons reside. The mouse auditory cortex contains multiple distinct areas that surround the primary auditory cortex (A1), including the anterior auditory field (AAF), the secondary auditory cortex (A2), and a dorsal posterior field (DP), among others. Although the specific functions of each auditory cortical field remain unresolved, different fields process sounds in distinct ways. For example, while A1 neurons often respond to individual tone frequencies (Mizrahi, Shalev, and Nelken 2014), A2 neurons are significantly more responsive to multi frequency sounds (Romero et al. 2020; Kline, Aponte, and Kato 2023) and AAF receptive fields are biased towards faster temporal structure (Linden et al. 2003).

In addition to their different roles in sound processing, different auditory cortical fields also have distinct connectivity to motor centers (Henschke, Price, and Pakan 2021; Tsukano et al. 2017, 2019; Gămănuţ et al. 2018). One important modulator of auditory cortex during locomotion is a long-range projection from the secondary motor cortex, which synapses locally in auditory cortex onto both inhibitory and excitatory cells (Nelson et al. 2013; Schneider, Nelson, and Mooney 2014; Schneider, Sundararajan, and Mooney 2018). Projections from the secondary motor cortex innervate much of the auditory cortex, but its connectivity to the dorsal auditory cortex is denser than in primary auditory cortex (Henschke, Price, and Pakan 2021).

These differences in function and connectivity suggest that different cortical fields might be differentially modulated by movement. However, most previous studies that report movement related modulation have either specifically targeted the primary auditory cortex (Audette et al. 2022); have not reported the auditory cortical field from which data were collected (Schneider, Nelson, and Mooney 2014); or solely used anatomical coordinates to identify cortical fields (Henschke, Price, and Pakan 2021), which is better accomplished through *in vivo* functional mapping (Romero et al. 2020; Narayanan et al. 2023). In addition, prior studies have at most investigated only two auditory cortical fields at a time. It therefore remains largely unknown whether the heterogeneity of movement-related modulation in mouse auditory cortex may be related to the specific cortical field from which neural activity was recorded.

Here, we used *in vivo* wide-field imaging to functionally map four distinct auditory cortical fields. We then quantified the movement-related modulation of single neurons in identified cortical fields using two-photon calcium imaging. Our findings reveal heterogeneity in the magnitude and direction of movement-related modulation across individual auditory cortical cells. Despite some subtle and significant differences among cortical fields, the average modulation driven by locomotion and other spontaneous movements was largely consistent across all areas that we sampled. We conclude that movement-related modulation is pervasive throughout the auditory cortex, and that despite different roles in sound processing and differing connectivity, both primary and non-primary auditory cortex exhibit macroscopically similar modulations by behavioral state.

## RESULTS

### Sound-evoked responses in auditory cortex L2/3 are diversely modulated by locomotion

To understand how auditory cortex activity is influenced by movement, we made large-scale optical recordings from excitatory neurons in L2/3 of primary and non-primary auditory cortex of awake, behaving mice. GCaMP6s was expressed in excitatory neurons using transgenic breeding strategies, followed by wide-field and cellular imaging in multiple auditory cortical fields in the same mice (see Methods). Head-fixed mice were acclimated to running on a non-motorized wheel for 7-10 days prior to imaging (Fig. 1A) (Schneider, Sundararajan, and Mooney 2018).

**Fig.1:**
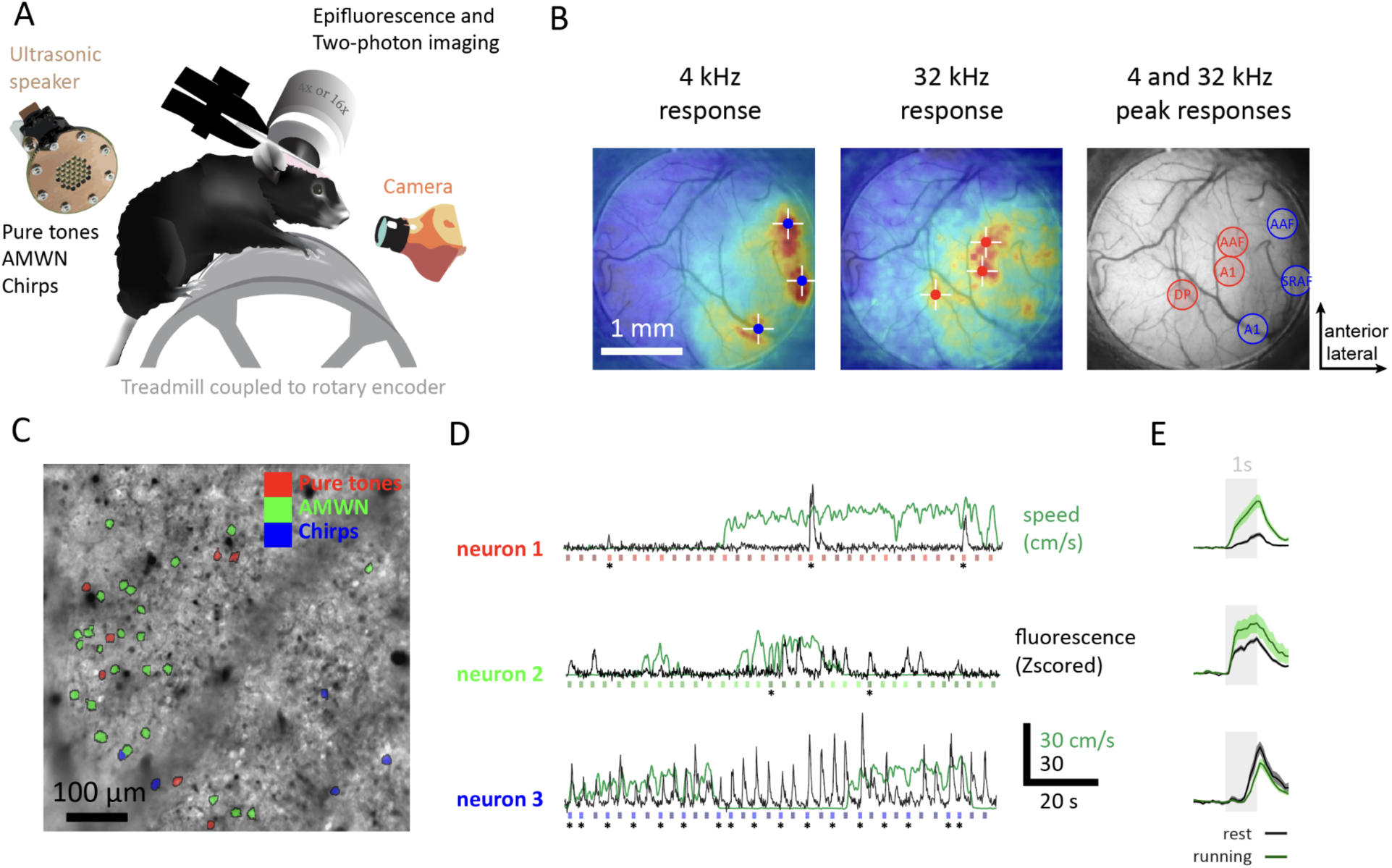
Mapping AC areas and cellular imaging of AC neurons during rest and locomotion across sound types. A. Schematic of the experimental setup used to map auditory cortex (AC) areas and conduct cellular imaging of AC neurons during rest and locomotion across different sound types. The setup involves a headfixed mouse running on a treadmill while sounds, such as pure tones, amplitude modulated white noise (AMWN), and chirps, are delivered through an ultrasonic speaker. High speed videos are recorded using a camera, and speed is measured through a rotary encoder attached to the treadmill. Auditory cortex images are acquired either through a 4X objective for epifluorescence AC mapping or a 16X objective for two-photon imaging through a 3 mm glass window. B. Epifluorescence auditory cortex mapping. Left and center, example epifluorence average sound responses to 4 and 32 kHz pure tones. White crossed show peaks. Right, peak 4 kHz and 32 kHz responses are represented by blue and red circles, respectively. C. Example two-photon field of view enhanced image, with sound responsive neurons being highlighted. The preferred sound type of each neuron is represented by colors in the legend. D. Example neuronal calcium and speed signals of 3 neurons. Calcium normalized fluorescence traces are represented in black and sound type, time and duration are represented by colored rectangles. Asterix represents preferred sound stimulus. Green line represents speed in cm/s. E. Peri-stimulus time histogram (PSTH) showcasing the example neurons individual preferred average and standard error sound responses during rest (black line) and running (green line) conditions. Sound timing is represented by a gray rectangle.

Using wide-field calcium imaging, we first identified four distinct auditory fields, including primary auditory cortex (A1), anterior auditory field (AAF), secondary auditory cortex (A2), and dorsal posterior area (DP) (Fig. 1B, Fig. 1S1,1S2) (Romero et al. 2020) (see Methods). We then used two-photon calcium imaging to measure sound-evoked responses from individual neurons in different auditory cortical fields (Fig. 1C). On each two-photon imaging day, one or more fields of view were imaged as mice rested and ran on the treadmill while hearing experimentally controlled sounds that were uncoupled from the mouse’s behavior (Fig. 1D). The sounds were drawn from three distinct categories: pure tones of varying frequency, amplitude-modulated white noise of varying modulation rates (AMWN), and up- or down-sweeping chirps. Overall, we recorded 1869 sound-responsive neurons (see Methods, Fig. 1E).

We first analyzed the sound tuning of individual auditory cortex cells while mice were at rest (Fig. 2A). Across the sound-responsive population (n=1869), the majority of neurons (75%) were responsive to one or more sound types during rest. Most neurons (57%) showed selectivity towards sounds from only one of the three categories, indicating a preference for specific types of sounds (Fig. 2B). Of the three categories, the majority of neurons were responsive to sounds in the AMWN category (27%), potentially because this category included more unique sound stimuli (9 AMWN frequencies versus 4 pure tones and 2 chirps) and therefore were more likely to have at least one sound within a neuron’s receptive field. A smaller subset of neurons (16%) were responsive to sounds in two different categories, while fewer than 2% of neurons were responsive to sounds in all three categories.

**Fig.2:**
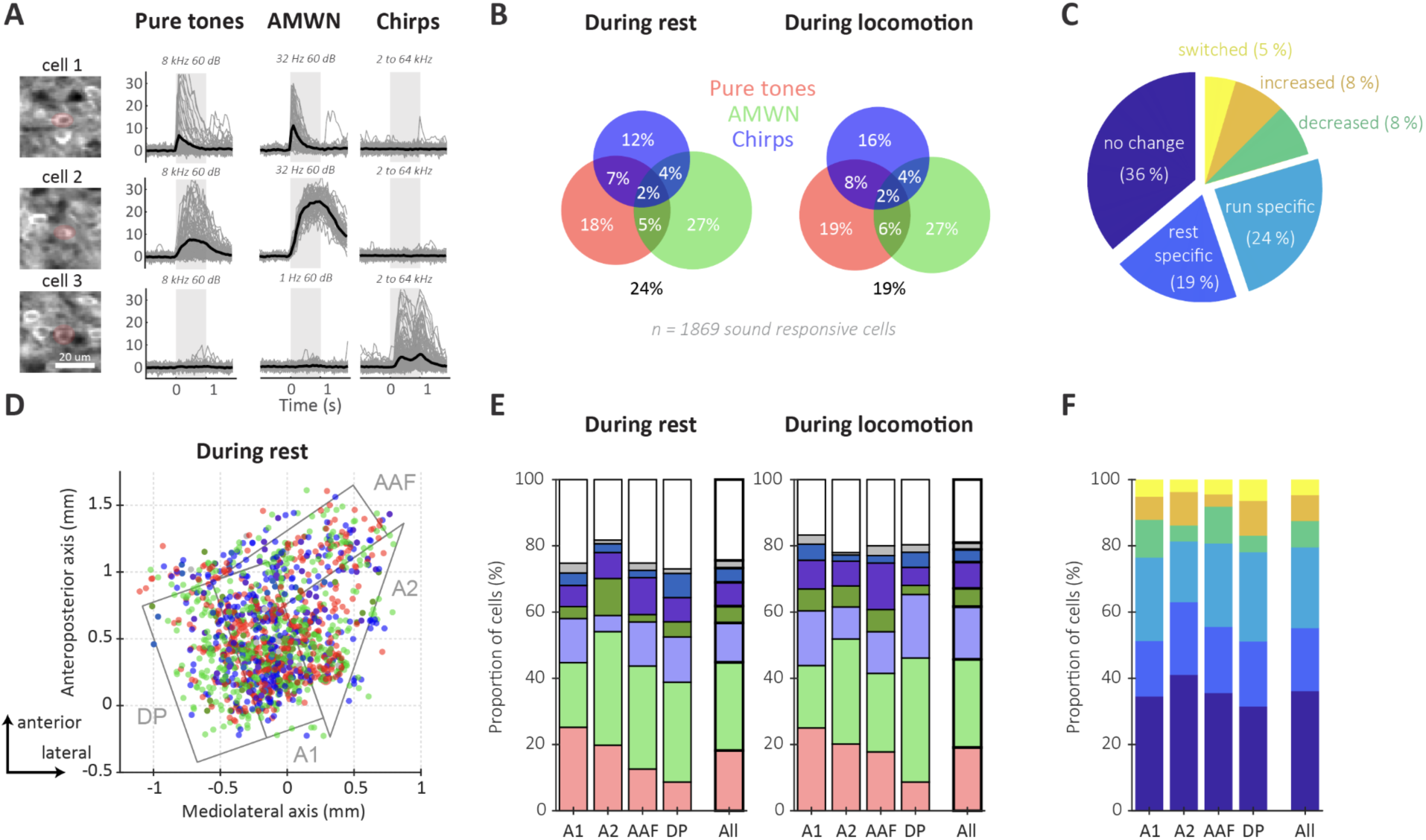
L2/3 neurons preferably respond to one type of sound and behavioral state but their preferences do not cluster spatially. A. Example sound responses. Left, Highlighted imaged layer 2/3 cells. Right, Fluorescence responses to sound (gray shade). Pure tone, amplitude modulated white noise (AMWN) and chirp responses are represented in consecutive columns. Black traces represent the average. Gray lines represent individual trials. B. Venn diagrams illustrating the proportions of neurons that respond to sounds during rest and locomotion for different sound types. C. Pie chart summarizing sound-type specificity changes between rest and locomotion states. The chart categorizes the neurons into four groups based on their responsiveness: those that show no difference in responsiveness, those that are exclusively responsive during rest or running conditions, and those that change their responsiveness during locomotion. The neurons in the last category can either increase or decrease the number of sound types they respond to or switch their selectivity to another sound type. D. Map of auditory cortex cells sound class preference shown as colored dots. The color of the dots uses the same color code as in B. Auditory fields are delineated in gray. The map is aligned to the 4 kHz A1 epifluorescence sound response. E. Sound type preference broken down by area. Color code is the same as in B. Chi-Square test revealed that the proportions of sound class preferences were significantly different during rest (p < 0.001), but not during locomotion (p > 0.05). F. Specificity change broken down by area. Color code is the same as in C.

We next asked how sound responses were influenced by locomotion. During locomotion, the proportions of neurons responsive to each sound category were similar to the resting condition, with 2% of neurons responsive to sounds in all three categories (Fig. 2B). 24% of neurons were not responsive to any sounds heard during rest but were responsive to at least one sound while the mouse was locomoting (e.g. run specific on Fig. 2C), indicating that some auditory cortex cells are selectively responsive only when a mouse is running. A smaller fraction of neurons (19%) were sound-responsive only when the mouse was at rest. 36% of neurons were responsive to sounds during both behavioral conditions and maintained their tuning to the same sound category across conditions. A small fraction of neurons became responsive to fewer categories during locomotion (8%), some became responsive to sounds in more categories (8%), and only 5% of neurons switched their responsiveness from one sound category to another. Taken together, these results show that locomotion leads to a slight increase in the number of sound responsive neurons in the auditory cortex and that neurons largely retain their sound-category responsiveness across periods of rest and locomotion.

### State-dependent sound responses are consistent across auditory cortical fields

The auditory cortex comprises several distinct fields that are thought to be part of a hierarchy of auditory processing (Wessinger et al. 2001; Sharpee, Atencio, and Schreiner 2011; Kline et al. 2021). Different fields receive different amounts of input from motor regions of the brain and might be influenced differentially by the behavioral state of the mouse (Henschke, Price, and Pakan 2021). Therefore, we next separated neurons based on their anatomical location within functionally defined auditory cortical fields (Fig. 2D), as assayed by wide-field calcium imaging (Romero et al. 2020; Liu et al. 2019).

For each mouse, we aligned wide-field imaging data to a stereotyped map of the cortical surface, (see Supplementary Fig. 1S1) and we overlaid all sound-responsive neurons onto a map which was color coded by the stimulus that drove the largest (and typically only) response (Fig. 2D). By plotting only the pure tone responsive cells based on their best frequency tuning, our two-photon data confirmed the expected tonotopic gradient of A1 (Fig. 1S2A,B). In contrast, when plotting cells tuned to AMWN, we observed no periodotopic gradient across the A1 tonotopic axis (Fig. 1S2C,D). During rest, we observed neurons responsive to all sound categories in each cortical field and we found that individual neurons throughout the auditory cortex tended to be tuned to sounds from just one sound category (Fig. 2D,E). To test this more definitively, we drew boundaries around approximate cortical fields and classified neurons as belonging to each field (Fig. 2D). During rest and locomotion, the fractions of neurons that were responsive to each sound category were significantly different across auditory cortical fields (Chi Square test, rest : p<0.001, locomotion: p<0.001; Fig. 2E). The largest fractions of neurons that responded only to tones were found in A1 (25 %), consistent with more primary sensory regions encoding individual sound frequencies (Mizrahi, Shalev, and Nelken 2014). The proportion of neurons responsive to AMWN was larger in non primary areas (A2= 34%,AAF= 31%, DP=30%) compared to A1 (19%), consistent with higher level cortical regions encoding more spectrally broad and complex sounds (Romero et al. 2020).

In all fields of the auditory cortex, more neurons were responsive to sounds during locomotion compared to rest (Fig. 2E). As observed across the broader auditory cortical population, only a small subset of neurons in each cortical field changed their tuning from one sound category to another when mice were locomoting compared to resting (Fig. 2F). Altogether, these results indicate that primary and non-primary cortical fields process sounds in subtly yet significantly different ways, but that locomotion-induced changes in category tuning of sound-evoked responses are largely consistent across auditory cortical fields.

### Locomotion-related changes in response magnitude are locally diverse and globally similar

In the analyses described above, we binarized neurons as either being responsive to, or unresponsive to, sounds during each of the two behavioral states (resting and running). We next more thoroughly quantified how the magnitudes of neural responses were influenced by locomotion. Individual auditory cortex neurons were diversely modulated by locomotion, with some neurons having larger responses, some neurons having largely stable responses, and other neurons having weaker responses during locomotion (Fig. 3A). Of the 1869 neurons that were sound responsive in at least one behavioral condition, most had larger neural responses during locomotion compared to rest, leading to average responses that were 15% larger during locomotion (average sound response R, R_rest_ = 0.45, R_locomotion_ = 0.57, Fig. 3A,B). To analyze responses at the level of individual neurons, we calculated a modulation index value for each neuron that described the ratio of responses during locomotion and rest (range: -1 to 1, see Methods) (Fig. 3C). The distribution of modulation index (MI) values skewed positive, indicating that on average, sound-evoked responses were stronger during locomotion compared to rest (MI = 0.07±0.01). We also scattered the magnitude of each neuron’s response during rest and running and fit a regression line to the data (Fig. 3D). While the average response was larger duringrunning compared to rest (see Fig. 3C), we noted that this derived primarily from a positive offset of the intercept accompanied by a slope that was less than 1, rather than a steeper slope of the regression line, suggesting that smaller sound responses are enhanced by locomotion while bigger responses are suppressed.

**Fig.3:**
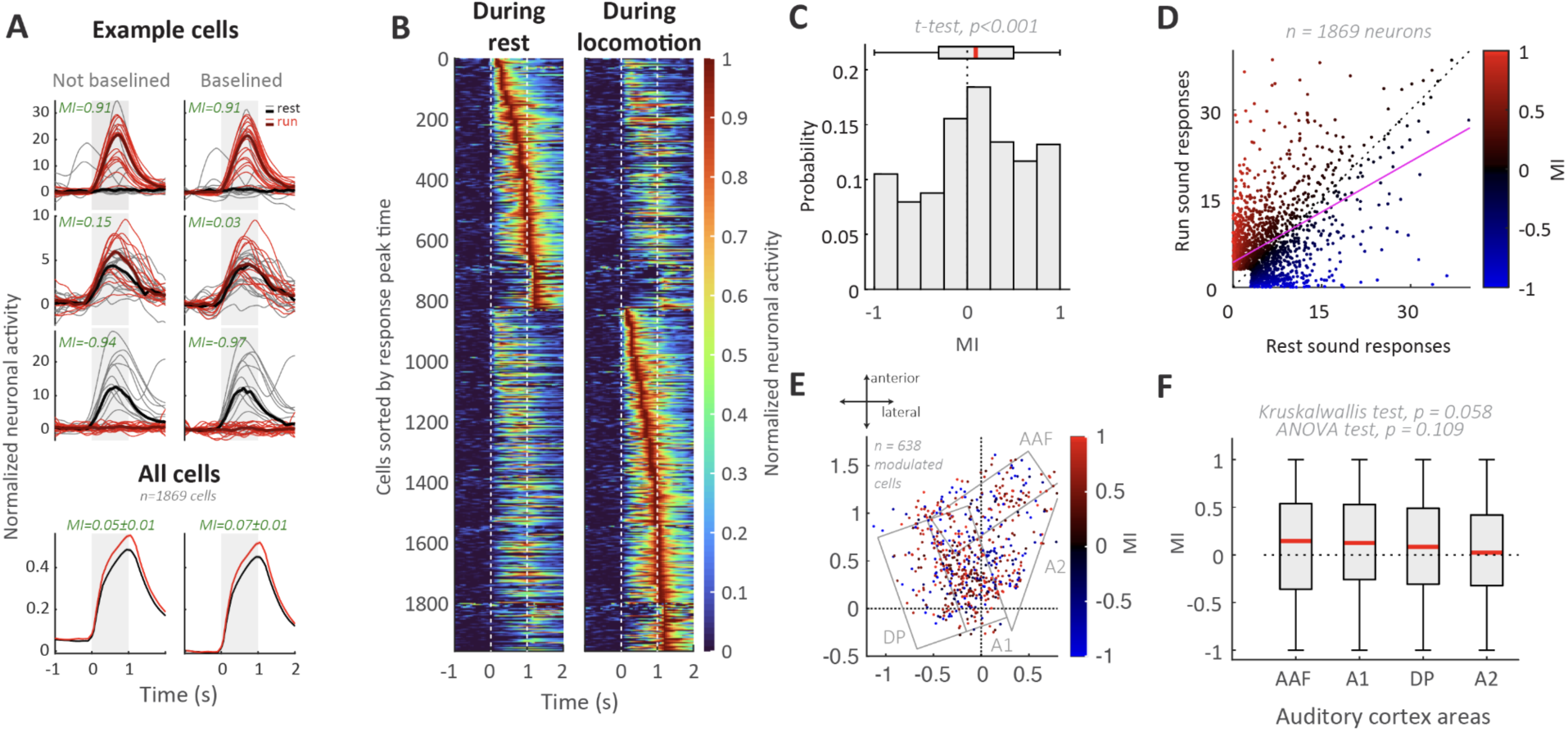
Running increases auditory cortex sound responses similarly across areas. A. Example (top rows) and average (bottom) sound responses in resting and running condition. Running modulation indexes (MI) are reported in green. Each sound lasted 1 second and is represented by a gray shade. B. Heatmap illustrating individual normalized sound responses during rest (left) and locomotion (right). Start and end times of sound are delineated by the white dotted lines. C. Distribution of modulation indexes with boxplot and red line indicating the median and quartiles of the distribution. T-test significance indicates a slightly positive population modulation index. D. Neuronal sound responses under rest and running conditions, with each dot representing a neuron and the color indicating its modulation index (MI). Small sound responses were excluded (see Methods). The dotted line represents the unity line, while the pink line shows the linear regression fit. E. Spatial organization of running modulated cells illustrated as MI map with each dot representing a significantly modulated cell. Significance was determined as sound responses in rest and running being significantly different. Grey lines delineate auditory cortex areas. F. Area-specific modulation indexes reported by boxplot (median and quartiles). Significance test reported in gray.

We next looked at the distribution of modulation index values across different auditory cortical fields. To visualize modulation across the cortical surface and across cortical fields, we overlaid all sound-responsive neurons onto a stereotyped map, color coded by their modulation index (Fig. 3E). Neurons that were strongly positively and strongly negatively modulated during locomotion appeared to be distributed rather uniformly across the broader auditory cortical surface (Fig. 3E). Dividing the auditory cortex into spatial bins and analyzing the mode, mean, or median of each bin did not show any spatial gradient (Fig. 3S2A-C). To test this more definitively, we drew boundaries around approximate cortical fields and measured the distribution of modulation index values for neurons within each field. We found that the average modulation index and range of modulation indices were similar across all cortical fields, with AAF and A1 tending to be more enhanced by locomotion compared to DP and A2 (Fig. 3F). Within A1, we found that modulation remained consistent across best frequency tuning ranges (Fig. 1S2E,F).

### Locomotion speed is encoded across auditory cortical fields

We find strong locomotion-related changes in neural activity that are heterogeneous across neurons and largely homogenous across cortical fields. But locomotion is not necessarily an all or nothing event, and mice can run fast, slow, and in between. In addition, other factors could also influence sound-evoked activity during behavior, such as how strongly a neuron responds to sounds during rest (Fig. 3D; Supplementary Fig. 3S1), the latency of a neuron’s peak response during sound playback (Fig. 3S1F), the cortical field in which a neuron resides (Fig. 3S1B), the depth within layer 2/3 at which a neuron resides (Fig. 3S1D), or the sound type to which a neuron is responsive (Fig. 3S1C). We reasoned that these attributes could all reflect aspects of a neuron’s function within the auditory cortical hierarchy and could presumably relate to whether or how a neuron’s sound-evoked response is influenced by locomotion.

To determine which factors most strongly contributed to a neuron’s locomotion-related modulation, we first segregated trials (i.e. sound events) based on the speed at which the mouse was running at the time of sound playback. We found that as the mouse ran faster, the modulation index increased significantly (Fig. 4A). We then calculated the effect size of trial speed and other variables (Fig. 4B), which revealed that running speed had the largest impact on neural activity, followed by the strength of a neuron’s response to sound during rest.

**Fig.4:**
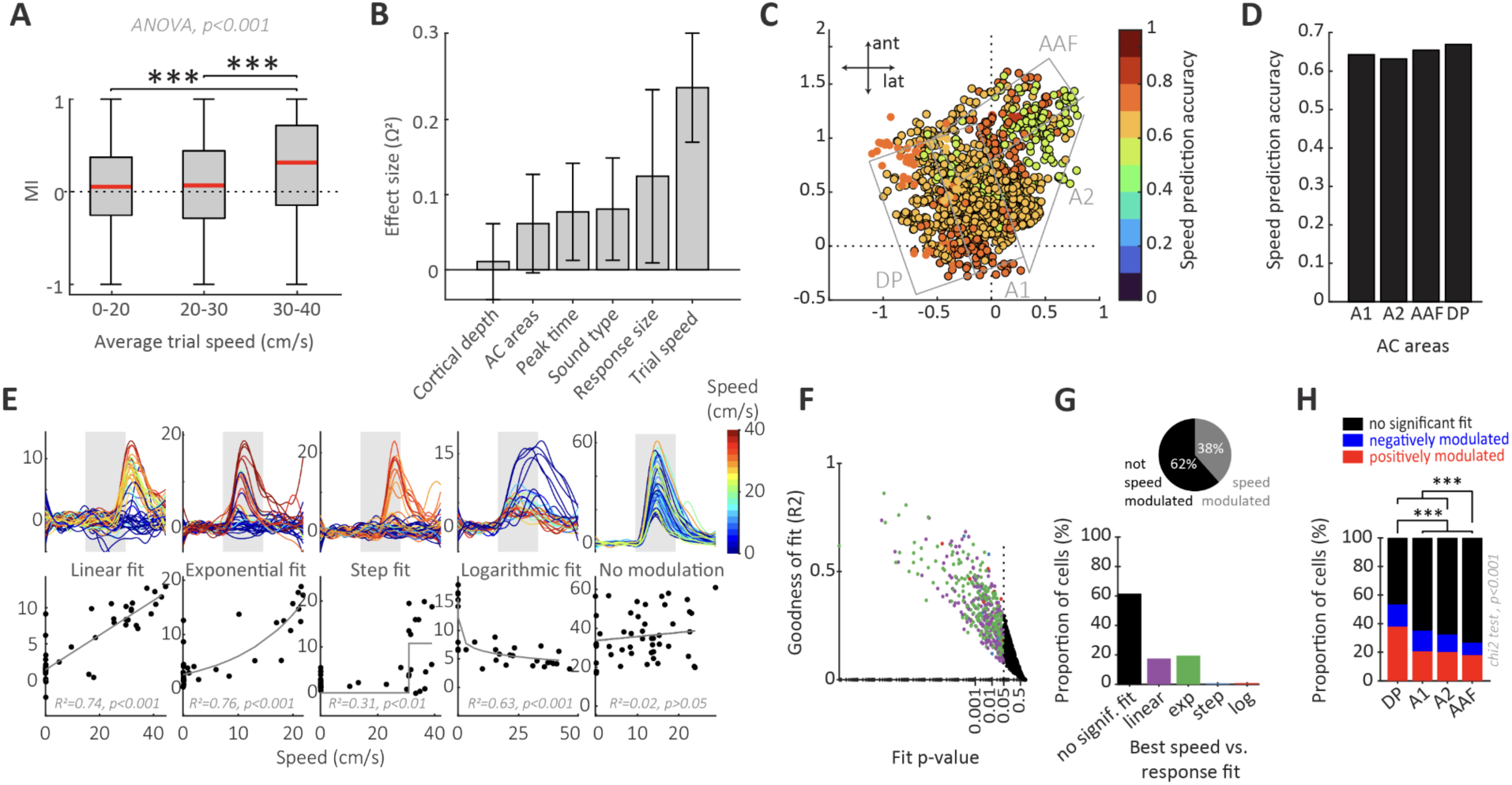
Auditory cortex contains speed-modulated neurons. A. Average trial speed relationship to modulation index (MI) shown as boxplots for various trial speed bins. Significance test reported in gray. B. Influence of recording variables on neuronal modulation index, as computed by the effect size (ω^2^,see Methods). Error bars represent 95% confidence intervals of the effect size. C. Running speed decoding accuracy map of the auditory cortex. Circles represent neurons and their colors, how accurate speed decoding was for that recording session, as shown in the color bar. The black edge on the circle indicates that decoding of the speed of this recording was significant (SVM permutation test, see Methods). Gray lines delineated auditory cortex areas. D. Bar plot of average speed decoding accuracy for individual areas of the auditory cortex. E. Example cells’ sound responses modulated diversely by speed. The top row shows sound responses, with each line representing a trial and the color representing the trial speed. Gray background box represents sound timing. The bottom row shows the relationship between speed and sound response size. Data are fitted by functions denoted in the legend. F. Scatter plot of all neurons (dots) showing the goodness of the fit (measured by R2), the fitting p-value, and the best fit shown by the color of the dots (legend in E). Significance alpha is symbolized by a dotted line at p=0.05. Black dots represent neurons with no significant fit. G. Bar plot quantifying the proportions of neurons’ best fits among linear, exponential (exp), step function (step), and logarithmic (log). The pie chart summarizes the proportion of speed modulated cells. H. Bar plot showing the proportion of speed-modulated cells across individual areas of the auditory cortex.

Given the importance of running speed on neural responses, we next attempted to decode the mouse’s running speed during sound playback from populations of simultaneously recorded neurons in each distinct auditory cortex field (Fig. 4C). For each recording session, we trained a linear support vector machine while holding out one of the trials, and then predicted the running speed on the held out trial (i.e. leave-one-out cross validation; see Methods). We considered decoding accurate if we could predict the running speed within 2 cm/s on a trial. We found that running speed could be decoded with around 65% accuracy from most recording sessions (Fig. 4C,D). While there was heterogeneity in decoding accuracy across sessions, this variability did not map onto distinct cortical regions; decoding accuracy was equivalent in A1, A2, AAF, and DP (Fig. 4D).

Given that running speed can be decoded at the population level, we next wanted to identify how running speed was encoded at the level of individual neurons. We identified individual neurons with responses that were enhanced during locomotion, suppressed during locomotion, or unmodulated. For each neuron, we fit four different models to its sound-evoked response magnitude at different running speeds (linear, exponential, step, and logarithmic; see Methods), and we identified the model that explained the most variance in neural responses across trials (Fig. 4E). Neurons for which no model could achieve a significant fit to the data were categorized as non-modulated (Fig. 4E,F,G). Across all of auditory cortex, we found that the majority of neurons (62%) were not significantly modulated by speed (Fig. 4G). Of the remaining 38% of neurons, the majority were best fit by either a linear or exponential model, with roughly equal proportions (Fig. 4G, p_linear_ = 16%; p_exp_ = 19%; p_step_ = 1%; p_log_ = 2%).

Lastly, we categorized neurons by cortical field and compared the fraction of neurons that were non-modulated, positively modulated, and negatively modulated by locomotion. All fields contained neurons that were both positively and negatively modulated (Fig. 4H), consistent with our earlier analyses (Fig. 3E). Area DP had significantly more locomotion-modulated neurons than did any other area, with AAF having the fewest number of locomotion-modulated neurons (Fig. 4H, DP = 51%, A1 = 35%, A2 = 32%, AAF = 27%). Our findings indicate that speed modulation is distributed throughout the auditory cortex. Although most individual neurons are not significantly modulated by locomotion, speed can nonetheless be reliably decoded from small populations of neurons throughout the auditory cortex.

### Auditory cortex neurons are modulated by multiple different movements

Although our analyses thus far have focused on one behavior, locomotion, we noted that mice also perform other behaviors while head restrained atop the treadmill. These behaviors include resting, grooming, and postural adjustments, as well as finer distinctions between different types of locomotion (e.g. walking and running). This provided an opportunity to determine whether the patterns of movement-based modulation we observe for running were similar across other types of behavior. Therefore, we asked whether these different behavioral states were reflected in distinct modulation patterns across the auditory cortex.

To begin, we hand-labeled video frames when sound playback occurred, and categorized each sound playback event as rest, running, walking, postural adjusting (“adjusting”) (Ramadan et al. 2021), sniffing, or grooming (see Methods) (Fig. 5A). For visualization purposes, we performed dimensionality reduction (UMAP) on individual video frames, allowing us to embed each frame as a single point in a low-dimensional space. This low-dimensional embedding revealed that distinct behaviors tend to be well separated using just 2 dimensions (Fig. 5A). We next used a pseudocolor map to label each frame based on the speed at which the mouse was moving (Fig. 5B). We found that resting, adjusting and grooming all occurred at speeds at or near 0 cm/s. Locomotion states largely formed a distinct island in UMAP space, and we segregated walking from running based on speed and gate. Across recording sessions, we found that mice engaged in all behaviors that we measured, albeit with different rates (Fig. 5C).

**Fig.5:**
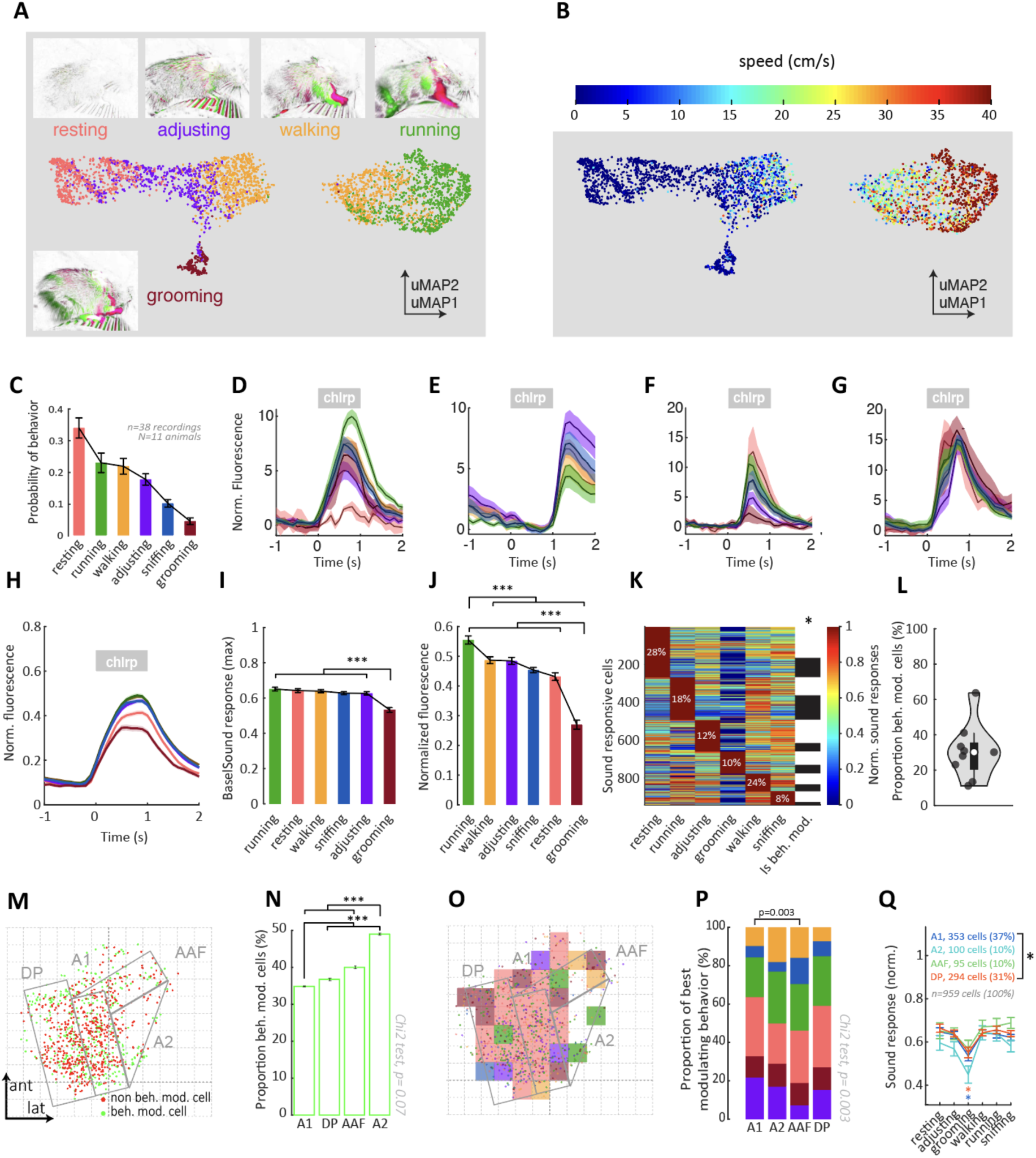
Auditory cortex sound response modulation depends on the type of movement. A. Example labeled session presented as a uMAP plot, where motion-frames are dots and color coded based on behavioral labels, such as resting, grooming, adjusting, walking, and running. Example motion-frames highlight motion in the previous frame in magenta and the next frame in green (see Methods). B. Similar to A, the uMAP plot displays motion-frames colored by running speed. C. Bar plot representing behavioral occurrences measured as the proportion of labeled frames. D to G. Example sound responses of neurons that are modulated differently by behaviors. H. Population average sound responses across different behaviors. I. Bar plot illustrating the size of sound responses (baseline subtracted) across different behaviors. Two-way analysis of variance revealed a significant main effect of areas (ANOVA, p<0.001) J. Bar plot depicting neuronal activity across different behaviors (sound responses non baseline subtracted), including both the baseline activity and the activity during sound responses. Two-way analysis of variance revealed a significant main effect of areas (ANOVA, p<0.001) K. Heat map displaying the activity of auditory cortex neurons under different conditions. Neurons are sorted based on their maximum sound response size, and the percentage of neurons with maximum responses is indicated in white. The last column of the heat map indicates whether neurons are differentially modulated by behaviors. The color represents the normalized neuronal response, as shown in the color bar. The color represents the normalized neuronal response, as shown in the color bar. The last column of the heat map indicates whether neurons are differentially modulated by behaviors (white = yes). L. Violin plot showing the proportion of behaviorally modulated neurons across multiple mice. M. Spatial distribution of cells that are differentially modulated by behaviors (green dots) compared to cells that are not influenced differentially by behaviors (red dots). The dashed lines indicate the reference point of A1 at 4 kHz, against which the maps are aligned. N. Bar plot quantifying the proportion of behaviorally modulated cells in each auditory cortex area. Bootstrapping analysis shows that the proportion of behaviorally modulated cells in A1 and A2 significantly differ from other areas (ANOVA, p<0.001, post hoc Tukey-Kramer with Bonferroni correction). Error bars represent 99% confidence intervals of bootstrapping distributions. O. Spatial distribution of behaviorally modulated cells depicted by dots. Dot colors represent the best sound response conditions. In the background, a density plot shows the most common best sound response condition in a square of 200 µm. Gray lines delineate the limits of the areas of the auditory cortex. P. Bar plot presenting the proportions of preferred behavioral conditions in each auditory cortex area. Significant difference between proportion of favorite modulation between areas was revealed by performing a Chi-Square test (p = 0.003). Q. Error bar plot showing normalized sound responses across different behaviors and auditory cortex areas. Two-way analysis of variance revealed a significant main effect of behaviors and areas (ANOVA, post hoc Tukey’s HSD with Bonferroni correction, p_Areas_<0.001, p_Behaviors_<0.001, p_Interaction_>0.05).

Next, we compared neural responses to sounds during each different behavior. We identified many individual neurons with responses that varied across behavioral states (Fig. 5D-F), and other neurons with responses that were largely invariant to state (Fig. 5G). Compared to rest, sound evoked responses and spontaneous fluorescence were on average enhanced during running, walking, postural adjusting, and sniffing, while neural responses were suppressed during grooming (Fig. 5H). Baseline-subtracted sound-evoked responses were significantly weaker during grooming than during all other behavioral states (Fig. 5I). When the baseline was not subtracted, we found that responses during running were significantly higher than all other behaviors and that responses during grooming were significantly lower (Fig. 5J). Across the population of sound-responsive neurons, distinct groups of neurons had their strongest sound evoked responses during different behaviors and for a third of the neurons, their activity for one behavior was significantly different than that measured during other behaviors (Fig. 5 K-L). Together, these findings reveal that auditory cortex is modulated by many behaviors, that not all behaviors modulate the auditory cortex equally, and that sound-evoked changes and baseline changes in activity both contribute to movement-related modulation in the auditory cortex.

Finally, we asked whether distinct auditory cortical fields were differentially modulated by behaviors other than locomotion. We plotted the spatial distribution of cells, color-coded by whether or not they were modulated by each behavior (Fig. 5M). Overall, we found that A1 had the lowest fraction of behavior-modulated neurons, while A2 had the highest fraction (Fig. 5N). Across the auditory cortical surface, we did not observe any clear dominant modulation by a particular movement (Fig. 5O, Fig. 5S1). However, A1 and AAF neurons differed significantly in the relative distributions of movements that modulated their activity (Fig. 5P). A2 tended to have weaker responses across most stimuli (Fig. 5Q). And across all areas, sound-evoked responses during grooming were weaker than during any other behavioral state (Fig. 5Q). These data together reveal that movement-related modulation in general is widespread throughout auditory cortical fields.

## DISCUSSION

Here we demonstrate that neurons throughout L2/3 of the mouse auditory cortex are modulated by locomotion and other behaviors, including walking, grooming, sniffing, and postural adjustments. At the level of single neurons, we find that movement-related modulation can be diverse and heterogeneous. But this heterogeneity is distributed nearly uniformly across auditory cortical fields, revealing a homogenous influence of behavioral state at a more macroscopic level. These observations provide important insights into how the auditory cortex integrates motor- and sound-related signals during behavior.

The mouse auditory cortex contains multiple distinct regions that can be identified using functional landmarks and tonotopic gradients. However, those regions are difficult to locate using anatomical coordinates (Narayanan et al. 2023; Romero et al. 2020). Furthermore, different auditory cortical fields have different responses to simple and complex sounds (Kline, Aponte, and Kato 2023; Linden et al. 2003) and different long-range connectivity (Henschke, Price, and Pakan 2021; Tsukano et al. 2017, 2019; Gămănuţ et al. 2018). Despite these differences and despite the diverse modulation phenotypes that we observed during locomotion, we found that each cortical field at a macroscopic level had largely similar movement-related modulation. This includes a slight enhancement of sound-evoked responses during movement, a stability of sound-category tuning during both locomotion and rest, reliable encoding of locomotion speed during periods of running, and modulation by multiple uncued behaviors.

In addition to these similarities, we identified significant differences across auditory cortical fields. First, we noted that the largest fractions of neurons that responded only to tones were found in A1 and A2 and the largest fraction of neurons responsive only to AM noise was observed in DP, AAF and A2 (Fig. 2E). These observations align with a hierarchical change in the complexity of tuning curves across cortical fields (Mizrahi, Shalev, and Nelken 2014; Romero et al. 2020). Second, we saw that DP had more speed-sensitive neurons than any other field, while AAF had fewer speed-sensitive neurons than any other field (Fig. 3H). These differences in modulation may reflect underlying connectivity differences across areas, particularly with respect to long range behavior-related inputs (Henschke, Price, and Pakan 2021). Finally, when we expanded our analysis to include other behaviors, we found that A2 had the highest fraction of movement modulated cells while A1 had the lowest fraction (Fig. 5N). Our findings largely complement previous observations of diverse and distributed movement-related modulation across the auditory cortex, including subtly different modulation patterns in primary versus dorsal auditory cortex (Henschke, Price, and Pakan 2021).

While locomotion has been the most commonly studied behavior for sensory modulation in mice, other behaviors are also known to influence sensory cortex. Electrophysiological recordings made during forelimb behaviors reveal a net suppression of sound-evoked responses in the auditory cortex, but also heterogeneous effects at the single-cell level, including many neurons that are enhanced or unaffected by movement (Rummell, Klee, and Sigurdsson 2016; Audette et al. 2022). In addition, movements including running, grooming, postural adjustments, and vocalizing all lead to similar changes in subthreshold membrane potential dynamics, consistent with our observation of similar changes in calcium responses across multiple different behaviors (Schneider, Nelson, and Mooney 2014). Together with these prior reports, our findings suggest that the modulation observed during locomotion is not categorically different from the modulation observed during other behaviors. This consistency across behaviors could indicate that movement-related modulation is not related to movement *per se*, but may instead be related to arousal (McGinley, David, and McCormick 2015). However, arousal and movement tend to have dissociable effects on sensory cortical activity (Vinck et al. 2015) and different movements can be decoded from neural activity throughout much of the dorsal cortical surface (Mimica et al. 2018; Musall et al. 2019; Stringer et al. 2019). Consistent with these ideas, here we also find that different populations of auditory cortical neurons are differently modulated by distinct behaviors (Fig. 5P). We argue that rich movement-related information is a feature of sensory cortical activity that likely subserves perception while animals interact with the world.

One interpretation of the widespread modulation of auditory cortex is that behavior-related signals may be ubiquitously important for processing both simple and complex sounds. It also remains possible that behavior-related signals play different roles in distinct cortical fields. For example, while motor-related signals might be purely modulatory in some areas, they may serve as a teaching signal in others, such as when learning to anticipate the acoustic consequences of action (Schneider, Sundararajan, and Mooney 2018). Alternatively, movement signals may help animals learn to produce appropriate behaviors in response to different sensory input (Znamenskiy and Zador 2013). Distinguishing among the many possible roles for movement-related modulation throughout the auditory cortex will be an important focus of future experiments.

Previous studies using electrophysiological measures (i.e.. action potentials or membrane potentials) to quantify neural activity have observed suppressed sound responses during locomotion compared to rest (Schneider, Nelson, and Mooney 2014; Rummell, Klee, and Sigurdsson 2016; Audette et al. 2022; Zhou et al. 2014; Bigelow et al. 2019). In contrast, experiments using calcium indicators have reported enhanced responses during locomotion, consistent with our current observations (Vivaldo et al. 2022; Henschke, Price, and Pakan 2021). These differences in the direction of modulation could stem from multiple factors, including sampling from different layers when monitoring calcium compared to electrophysiology; longer temporal analysis windows for calcium indicators; and a non-linear relationship between action potentials and calcium levels. We also note that there are some similarities between experiments using calcium indicators and those using electrical methods. Both experimental techniques reveal a rich heterogeneity of movement-related modulation. And when we fit a regression line to compare sound-evoked responses during running and resting, we observed a slope of less than one, consistent with the suppression observed using electrophysiological recordings (Fig. 3D). Uncovering why different physiological recording methods reveal different distributions of modulation directions and magnitudes will be an important avenue of future investigation.

Finally, we note that the current experiments focus solely on excitatory neurons in L2/3. While this population has been previously shown to be strongly influenced by behavior (Audette et al. 2022; Keller, Bonhoeffer, and Hübener 2012), further differences in motor-related modulation might arise when sampling across cortical layers. Indeed, motor-related signals tend to be stronger in deep cortical layers compared to superficial layers (Audette et al. 2022). Moreover, excitatory and inhibitory cells are also differently influenced by behavior (Schneider, Nelson, and Mooney 2014) and signals related to violations from sensory expectations during behavior have recently been found to be concentrated in specific genetically defined populations of neurons in L2/3 (O’Toole, Oyibo, and Keller 2022). Future investigations with layer- and cell-type specificity may reveal more nuanced distinctions in how movement modulates different auditory cortical fields.

## SUPPLEMENTARY FIGURES

**Fig.1S1:**
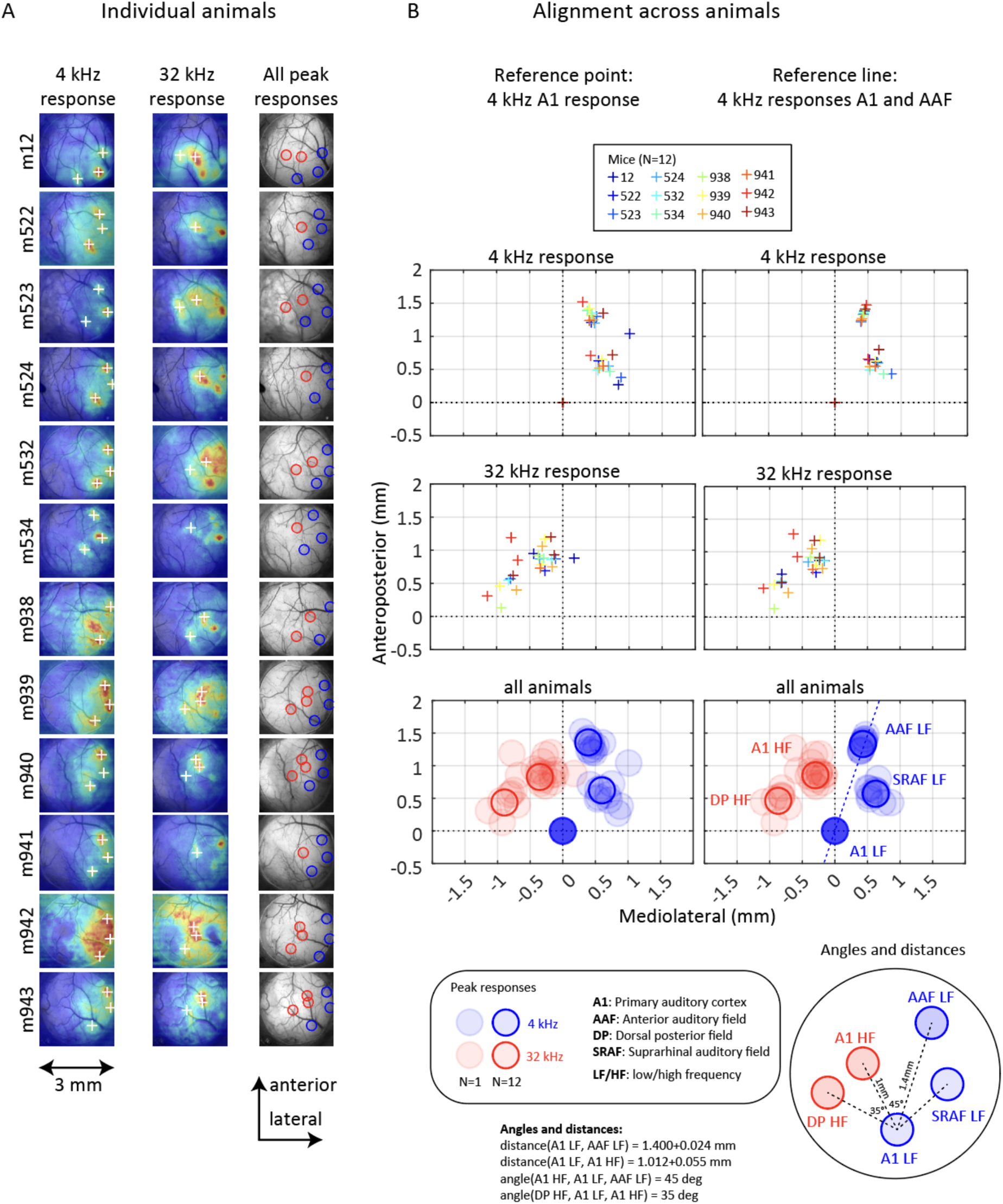
Auditory cortex mapping in individuals and across animals. A. Individual 4 kHz and 32 kHz responses of each animal, as well as the corresponding AC map. The legend for this figure is the same as in Figure 1. B. Two alignment strategies across animals using a point (A1 4 kHz) as reference (left) or a line (A1 4 kHz to AAF 4 kHz). In the first and second rows, plus signs indicate the peak responses of each animal after alignment. The third row of the figure shows the maps aligned across animals using the two different reference strategies, along with the average location of the area LF and HF reference points (low transparency circles). The figure also reports the average angles and distances between the landmarks.

**Fig.1S2:**
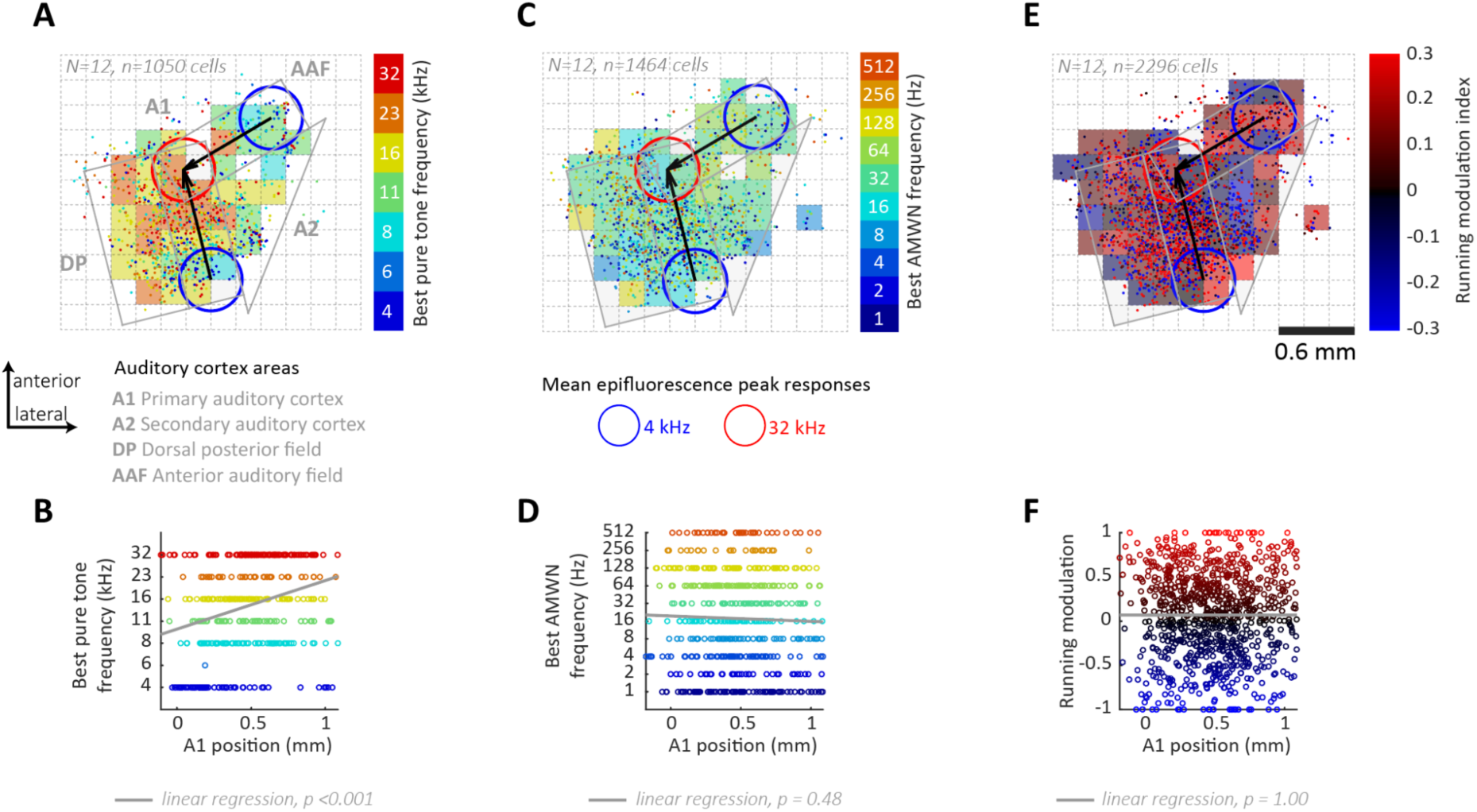
Best pure tone frequency of auditory cortex cells organize in tonotopic gradients. A. This pure tone best frequency map shows the location of neurons (dots) that respond to pure tones across 12 animals, with the color of each dot representing the neuron’s best frequency as identified in the color bar. AC was binned in 200 um tiles, of which color represents the average best frequency of the neurons. The gray lines on the map delimit the areas of the auditory cortex (AC) extrapolated from mean epifluorescence peak responses to 4 and 32 kHz (blue and red circles). The map is aligned to the primary auditory cortex 4 kHz responses, with the tonotopic gradients of A1 and AAF indicated by black arrows. B. The tonotopic gradient along the primary auditory cortex (A1) is described in this graph, which plots the position of cells along the A1 axis (indicated by the black arrow on A) against their corresponding best pure tone frequency. A linear regression model (shown in gray) was fitted to the data, revealing a significant relationship between A1 position and the best pure tone frequency (p < 0.001). C. This best frequency map shows the location of neurons that respond to amplitude modulated white noise (AMWN), with the color of each dot representing the neuron’s best frequency as identified in the color bar. The map legends are similar to panel A. D. Absence of periodotopic gradient along the A1 axis is revealed by a lack of a significant relationship between the A1 position and best AMWN frequency. E & F. Running modulation index map depicts all neurons colored by their modulation index. Neurons are organized in a salt-and-pepper fashion. Lack of significant relationship between A1 position and running modulation index.

**Fig.3S1:**
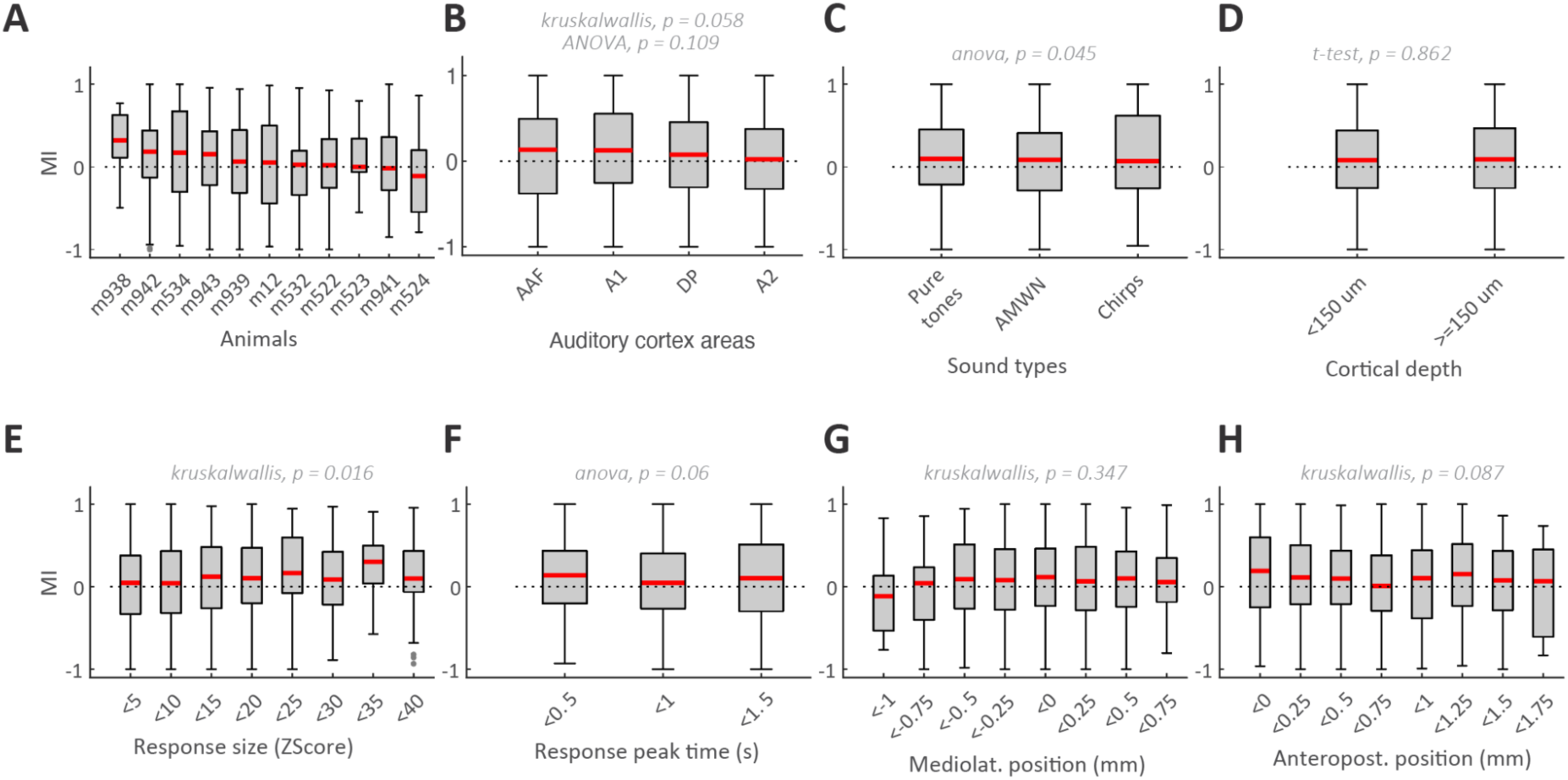
Only session average speed affects running modulation index. A, Boxplot showing median (red line) and quartile (grey box) modulation index distributions in individual animals. B to H, Distribution of cell modulation indexes binned in groups according to auditory cortex areas (B), sound type preference (C), cortical depth (D), response size (E), response peak time (F), mediolateral (G) and anteroposterior (H) positions. Statistical comparisons are reported in grey.

**Fig.3S2:**
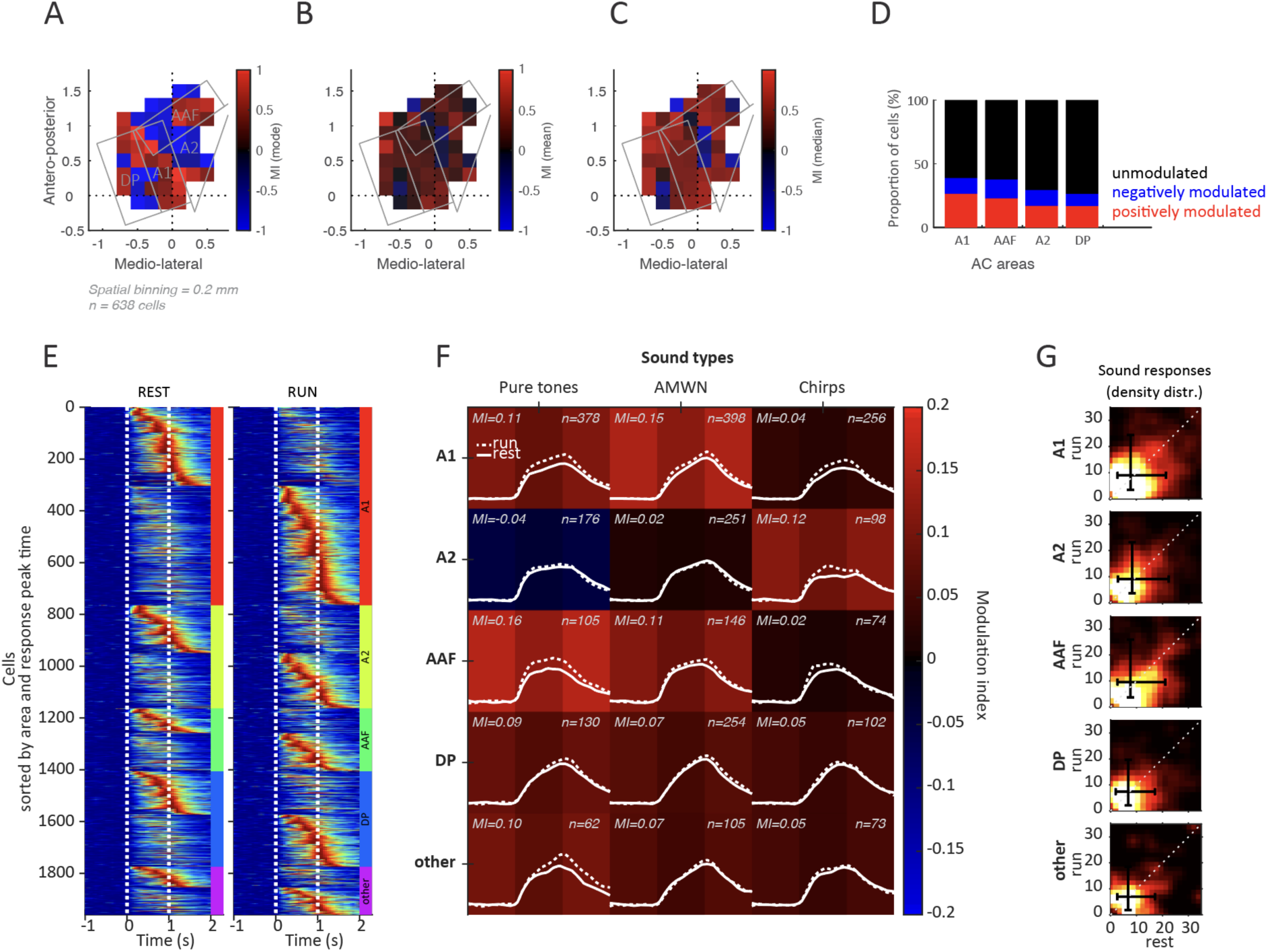
Running modulates sound responses similarly across areas and sound types. A,B,C. Spatial organization of modulation of AC modulated cells. Auditory cortex maps are binned in 0.2 mm bins with a minimum of 5 modulated neurons per bin. Gray lines represent the limits of AC areas. The color of each bin represents the mode (A), the average (B), or the median (c) modulation index across modulated neurons. D. Proportion of positively, negatively, and unmodulated neurons across auditory cortex areas. E. Heatmaps of sound responses by areas. Cells are sorted by areas and then by sound responses peak time. Dotted lines delineate the start and end of the sound. F. Summary matrix of average sound responses and modulation indexes by area and sound type. Background color represents the average modulation index. G. Density distribution plots showing rest versus running sound responses across different areas, with black error bars indicating the median and quartiles of each distribution.

**Fig.5S1:**
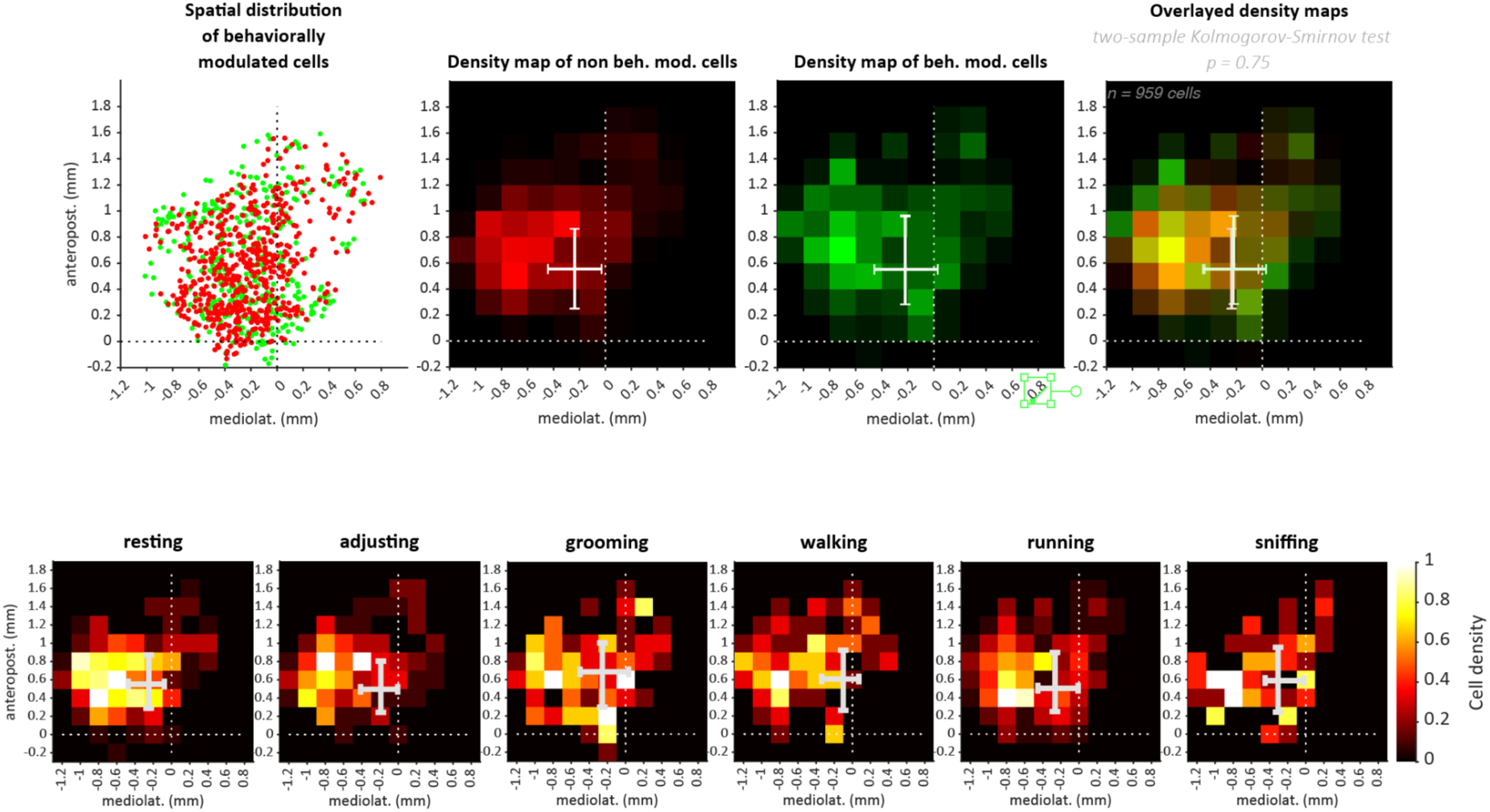
Spatial analysis of sound responses modulation by behaviors. A. Spatial distribution of cells that are modulated differentially by behaviors (green dots) as well as cells that are not influenced differentially by behaviors (red dots). The dashed lines indicate the reference point of A1 at 4 kHz, against which the maps are aligned. B. Density maps displaying the distribution of non-behaviorally modulated cells (red) and behaviorally modulated cells (green). Right panel depicts overlaid density maps allowing visualization of combined distributions. Error bars represent horizontal and vertical quartiles and medians of the spatial distribution of neurons. C. Density maps of cells modulated by distinct movements.

## METHODS

### Animals

Experiments were performed on 12 Camk2-Cre +/+ Ai148D +/+ (B6.Cg-Tg(Camk2a-cre)T29-1Stl/J;B6.Cg-Igs7tm148.1(tetO-GCaMP6f,CAG-tTA2)Hze/J, Jackson Laboratory, http://www.jax.org, RRID IMSR _JAX: 005359 and 030328) male mice (see table). Experimental animals were obtained by breeding homozygous colonies. Breeding cages were fed a doxycycline food diet for 3 weeks (625 mg/kg, ENVIGO, Teklad, TD.05125). This diet halts the Cre recombinase expression until 21 days of age, avoiding epileptic-like seizure activity in adults. Offsprings’ ear biopsies were genotyped by Transnetyx (Cordova, TN) for Camk2-Cre and Ai148D transgenes (Transnetyx probes for reference lgs7-1WT, eGFP, CRE). Positive offsprings were kept in grouped cages with regular food and water available ad libitum and kept on a reverse day-night cycle (12 hours day, 12 hours night). All procedures were carried out in accordance with New York University’s Animal Use and Welfare Committee. One mouse was excluded from the analysis because no sound-responsive cells were detected.

| Mouse ID | Number of responsive neurons | Number of recorded neurons | Percentage responsive (%) |
| --- | --- | --- | --- |
| Average | 222 | 6258 | 4.4 |
| 12 | 147 | 9787 | 1.5 |
| 522 | 366 | 9458 | 3.9 |
| 523 | 64 | 503 | 12.7 |
| 524 | 105 | 3038 | 3.5 |
| 532 | 362 | 11850 | 3.1 |
| 534 | 445 | 7186 | 6.2 |
| 938 | 47 | 1503 | 3.1 |
| 939 | 174 | 7480 | 2.3 |
| 941 | 152 | 5887 | 2.6 |
| 942 | 394 | 7896 | 5 |
| 943 | 189 | 4250 | 4.4 |

### Preparation of cranial windows

Glass coverslips were cleaned and stored in 70 ethanol distilled water solution. Dry 4 mm and a pair of 3 mm coverslips (#1 thickness, Warner Instruments) were glued using a transparent, UV-cured adhesive (Norlan Optical adhesive 61). On surgery day, cranial windows were rinsed with sterile saline (Argyle Sterile Saline 0.9 or Addipak Sterile Saline).

### Surgical procedure

Animals were anesthetized with isoflurane in oxygen (3 induction; 1.5–2 maintenance) and placed in a stereotaxic holder with non-rupture, zygoma ear cups (Kopf Instruments, model 963 and 1721) with a heating pad to maintain and monitor body temperature (Harvard Apparatus). The scalp was disinfected with 70 ethanol and Betadine. Mice were injected subcutaneously with an analgesic (0.1 mg/kg, Meloxicam ER, ZooPharm) and 150 uL local anesthetic under the scalp (0.25 bupivacaine hydrochloride saline solution, Sigma-Aldrich B5274-1G). Eyes were covered with Vaseline. After removing the scalp, the skull was dried and polished to remove the periosteum allowing better adhesion of the headpost. The dorsal part of the skull was covered with tissue adhesive (Vetbond, 3M), and a Y-shaped headpost (1.5 mm thickness, 120° angle, 3×17 mm branches, sendcutsend.com) was aligned to the midline and glued. The headpost was cemented to the dorsal skull using Metabond dental cement (C & B, Metabond) and the area over the skull was covered with a thin layer of transparent dental acrylic (Lang Dental). Once the dental cement was cured, the ear cups were removed to allow access to the auditory cortex (∼2mm diameter, -2.5mm posterior, 4.2 mm left from bregma), and the headbar was clamped to stabilize the mouse head. Temporalis muscle insertion on the temporal ridge was resected and glued down. The parietal bone was dried and cleaned, and a 3 mm circular craniotomy was drilled using a dental drill (Foredom, Model 1474) and micro drill bit (0.10 mm head diameter, GRAINGER, Kyocera, 07089). The posterior edge of the craniotomy borders the lambdoid suture, while the ventral edge is positioned below the squamosal suture located above the posterior end zygomatic arch. The coverslip was placed into the craniotomy with the help of a 3D manipulator-held wooden stick and hermetically sealed with Vetbond and dental cement applied between the surrounding skull and glass.

### Behavioral setup/Apparatus

Mice were trained for 7-10 days to run head-fixed on a 3D-printed wheel before imaging experiments. Head-fixation was achieved using a plate clamp (standa.it, 4PC69). The wheel was designed using Sketchup Free (https://app.sketchup.com/app, 40 mm diameter, 60 mm rod length, 1 mm rod diameters, 3 mm rod spacing) based on the previously published design (Villette et al. 2017) and affixed to a rotary encoder (https://www.usdigital.com/, H5-1000-NE-S). Sounds were played through an electrostatic ultrasonic speaker, amplified (TDT Tucker-Davis Technologies, ES1, and ED1), and controlled by a sound card (RME, FireFace UCX). The speaker was placed at about 10 cm from the right ear of the mouse (contralateral ear). Videos were acquired using IR zoom lenses (Xenocam, 9-22mm, 1.4 f) mounted on Sony CCD Camera (Amazon, AKK CA20 600TVL) and IR illumination was provided by infrared light sources (Phenas, Home 48-led CCTV IR lamp), and. Sound stimulation, video recording, and wheel position were controlled or recorded using custom-written code (MATLAB). Sound stimuli were played using Matlab Psychophysics Toolbox Version 3 (PTB-3) sound library (PsychPortAudio) (Brainard 1997).

### Auditory stimulation protocols for mapping and cellular imaging

All sounds were generated at 192 kHz using Matlab scripts.

#### Epifluorescence mapping auditory stimulations

Tonotopic mapping was achieved by presenting 200 ms pure tones (4, 8, 16, and 32 kHz at 80 dB) separated by 3 s and played pseudo-randomly such that each frequency was played 15-20 times. Mice were awake and head-fixed and free to move on the wheel. Stimuli were calibrated before recording using an ultrasonic acoustic microphone (Avisoft-Bioacoustics, CM16/CMP).

#### Cellular imaging auditory stimulations

A total of 38 sounds were played while head-fixed mice were free to run on the wheel. Sounds included 3 sound types: 7 pure tones (PT) (4 to 32 kHz spaced by 0.5 octaves), 9 amplitude modulated white noise (AMWN) sounds (modulation frequencies: 2, 4, 8, 16, 32, 64, 128, 256, 512 Hz) and 2 chirps (4 to 32 kHz and 32 to 4 kHz). PT and AMWN were played at 40 and 60 dB. All sounds were 1 s long with a rising and falling ramp of 5 ms. All sounds were presented pseudo randomly such that each sound was played 15 times in rest and running conditions.

### Calcium imaging acquisition

#### Widefield epifluorescence image acquisition for auditory cortex mapping

Widefield epifluorescence images were acquired with a 4x objective (Olympus, NA 0.10 Plan Acromat) focused 400 um below the cortical surface. A blue light-emitting diode (Thorlabs, M470L3, 470 nm, 650 mW) excited a ∼3 mm diameter area of cortex through a filter cube (Thorlabs, DFM1T1) containing an excitation filter (Thorlabs, MF469-35), a dichroic mirror (Thorlabs, MD498) and an emission filter (Thorlabs, MF525-39). Green fluorescence was captured at 30 Hz with a 16-bit camera (sCMOS, pco.edge 3.1). The sample was illuminated only during the acquisition to reduce bleaching (trial length 2 s). 384×512 pixel images were acquired using a Matlab custom GUI and Matlab image acquisition toolbox. A blood vessel map picture was taken at the pial surface as a reference image for targeting two-photon recordings.

#### Cellular imaging using two-photon microscopy

We used a resonant scanning two-photon microscope (Neurolabware, Los Angeles, CA) focused 150-200 um below the pial surface to image GCaMP fluorescence changes during behavior. Excitation was provided by a femtosecond pulsed laser (SpectraPhysics, Insight X3) tuned to 940 nm. Illumination power was controlled by Pockels cells (Conoptics, 350-80-02) and the beam size by a beam expander. The beam then passed through 8 kHz Galvo-Resonant Scanner (Cambridge Technology, 6215H galvo scanner and CRS 8K resonant scanner) and 16x/0.8NA water-immersion objective (Nikon, MRP07220) to form a 512×796 pixel field of view (∼1 mm^2^). The objective was rotated between 45-60° off the vertical axis until perpendicular to the cranial window to obtain images of the auditory cortex while maintaining the mouse head position straight. Two-plane imaging was achieved using an electrically tunable lens (Optotune). The frame rate was 14.49 Hz or 7.5 Hz for single- and two-plane imaging, respectively. Emission photons were filtered by photon collection optics (Semrock dichroics, 750 SWP, 562nm LWP; emission filter, 510/84 bandpass) and were detected by a GaAsP photomultiplier tube (Hamamatsu, H11706-40 MOD2). Images were written to disk at 1x,1.2x, or 1.4x digital magnification using an acquisition software (Scanbox software, Neurolabware).

### Image and data processing

#### Widefield epifluorescence for auditory cortex mapping

To localize the peak fluorescence responses to 4 kHz and 32 kHz pure tones on the cortex below the cranial window, images were processed using Matlab scripts. First, images were cropped only to include the 3 mm diameter cranial window view and resized 300×300 pixels to reduce processing time. Then, movies were averaged across 15-20 trials, baselined and Z-scored by the standard deviation for each pixel of the baseline period across all stimuli (0.2 s before stimulus onset). The response image for each stimulus was defined as the average across frames occurring during the response window (0.1 s to 0.3 s after stimulus onset). Fluorescence response peaks were detected on a smoothed response image (Gaussian filter radius = 100 um) using local maxima bigger than 25% of the tallest peak.

#### Auditory cortex areas definition

To identify auditory cortex fields (A1, primary auditory area, AAF, anterior auditory field, A2, secondary auditory areas, DP, dorsal posterior field), fluorescence responses to 4 kHz and 32 kHz pure tones were used as landmarks (Fig. 1S1). A1 was defined as a rectangle of 500 µm width spanning in length from the most posterior 4 kHz response to the most medial anterior 32 kHz response. AAF was defined as a rectangle of 500 µm width spanning from the most anterior 4 kHz responses to the most medial anterior 32 kHz response. DP was defined as a rectangle medial to A1 with a width of 600 µm. A2 was defined as the triangular area between A1 and AAF.

#### Two-photon microscopy

Two-photon calcium signals from auditory cortical neurons were extracted from imaging recordings using the toolbox Suite2p (Pachitariu et al. 2017). Briefly, suite2p corrected movement artifacts between frames with a non-rigid registration. Then, regions of interest (ROI) and neuropil areas (an annular ring surrounding the ROI) were identified. Then, calcium signals were extracted from soma and neuropil by spatially averaging pixel intensities across those regions for each ROI. Finally, ROIs were identified as somas during a manual curation step using the Suite2p user interface. ROIs were selected based on soma-like morphological features visible on the processed version of the fluorescence image (enhanced image). Each neuropil signal was then subtracted from the raw calcium traces to limit neuropil contamination. Neuronal signals were then baselined by subtracting the mode over a 60-second sliding window. Baselined traces were then normalized by the estimated noise for each cell. The noise distribution for each cell was defined as the distribution of negative values from the baselined traces and the same negative values multiplied by -1. The noise estimate for each cell was the standard deviation of that “symmetrized” noise distribution. For neuronal response computations, peristimulus histograms for each stimulus were baselined by the average of the 1 s pre-stimulus baseline period. The neuronal response of a neuron is reported as the peak value over a response window (0 to 1.5 s after stimulus onset) in units of standard deviations of the noise distribution of that neuron, referred to as “Normalized fluorescence” in the manuscript.

### Cell selection

Sound-responsive cells were defined by two criteria: (1) Significant statistical difference (Wilcoxon signed-rank test) between trial values from the baseline period (1s pre-stimulus) and at least one response window (0 to 0.5, 0.5 to 1, 1 to 1.5 s after stimulus onset). Using three response windows allowed to include onset and offset responses.. Response magnitude was calculated as the difference between the baseline period and the best response window. The minimum number of trials was set to 5 trials. Cells with lower number of repetitions were discarded. (2) Average response size above 3 standard deviations of the noise distribution of each neuron (see Image and data processing/Two-photon imaging).

### Data analysis

#### Alignment of auditory cortex maps across mice

To align the epifluorescence imaging maps of the auditory cortex across mice, we employed a two-step process that involved translation and rotation. First, we aligned all the maps to the 4 kHz frequency A1 epifluorescence sound response peaks. Next, we rotated individual maps such that the line joining the A1 and AAF 4 kHz epifluorescence response peaks would be at 70 degrees relative to the abscissa (Fig. 1S1B).

#### Modulation index computation

We calculated the modulation index (MI) as MI = (R_run_-R_rest_)/(R_run_+R_rest_), where R represents the preferred sound responses during rest (R_rest_) or run (R_run_). (Fig. 3)

#### Sound responses speed modulation analysis

##### Effect size

In Figure 4B, we compared how different recording parameters affect the modulation index. To do this in a standardized way, we used a statistical measure called effect size. Since our data had more than two parameters, we used a type of effect size called omega squared that estimates the variance due to an effect and dividing it by the estimated total variance, using the following formula: T ω² = (SS_between_ - (df_between_ * (SS_within_ / df_within_))) / (SS_total_ + (SS_within_ / df_withi_n))), where SS stands for sum of squares and df stands for degrees of freedom.

##### Decoding running speed from sound-evoked responses with support vector machines (SVM)

To estimate how much running speed information is carried by the sound-evoked neural responses of each recording, we employed the following decoding strategy. For each recording session, we included the sound responses elicited by all sounds, regardless of the preferred sound response of each neuron. This approach resulted in a matrix of dimensions ’number of neurons’ by ’number of trials’ and a vector containing the trial running speeds, computed as the average running speed during the sound presentation. Running speeds were binned into intervals of 2 cm/s, ranging from 0 to 40 cm/s.

We trained a support vector machine with a linear kernel to predict running speed based on the sound responses. As some speeds were more represented than others, we chose the ove-vs-one training strategy which better handles imbalanced datasets. To evaluate the performance of our decoding model, we employed leave-one-out cross-validation. Specifically, for each trial, we trained the SVM on the remaining trials and tested its prediction accuracy on the left-out trial. The accuracy of prediction was computed as the fraction of correct speed bin predictions when compared to the true speed bins. The regularization parameter C was 0.022.

To evaluate our results’ significance, we conducted a permutation test by reshuffling our data 100 times. In each reshuffle, we randomly altered the relationship between sound responses and running speeds, followed by training the SVM using the leave-one-out cross-validation approach as previously described, and assessed the prediction accuracy on the excluded trial.

Subsequently, we established the chance level accuracy by taking the average from the distribution of prediction accuracies across all reshuffles. We then contrasted the prediction accuracy derived from the original data with this distribution of chance level accuracies using a t-test, keeping the significance level threshold at 0.05. This procedure was replicated for all recorded sessions, and the resultant session accuracies were illustrated in Fig. 4A. The analysis was performed in MATLAB (MathWorks) using the function *fitecoc* from the Statistics and Machine Learning Toolbox.

##### Modeling the relationship between running speed and sound responses

To analyze the modulation of sound responses by speed, we applied different models to the data depicting the relationship between speed and sound responses for each neuron. We chose 4 fitting functions based considering the data’s visual pattern: linear, step-function, exponential and logarithmic (see Fig. 5C for examples). Following the fitting the observed values (y), we obtained the predicted values from the model (yfit) and proceeded to assess the quality of the fit using two metrics: the coefficient of determination (R²) and the p-value (Fig. 4D).

To calculate R², we first computed the residuals by subtracting the predicted values from the observed values (yresid = y -yfit). Then, we determined the sum of squared residuals (SSresid) by summing the squared values of the residuals. Additionally, we calculated the total sum of squares (SStotal), which represents the variability of the observed values around their mean. SStotal was obtained by multiplying the variance of the observed values by the length of the data minus one. The coefficient of determination (R²) was then calculated as 1 minus the ratio of SSresid to SStotal (R² = 1 - SSresid/SStotal). R² ranges from 0 to 1, where higher values indicate a better fit of the model to the data. To further assess the statistical significance of the fit, we calculated the p-value. The p-value was obtained using the F-distribution cumulative distribution function (CDF) with the F-statistic, which was computed as (R²/(1-R²)) multiplied by the ratio of the degrees of freedom (df) to 1. The degrees of freedom (df) were determined as the length of the data minus 2. Finally, the p-value was calculated as 1 minus the cumulative distribution function (CDF) of the F-distribution, using the F-statistic, 1 as the numerator degrees of freedom, and df as the denominator degrees of freedom.

Finally, for each neuron, the best fit was determined as the fit with the biggest coefficient of determination (R²) and a significance level of the fit below 0.05 (Fig. 4E).

#### Video processing and behavioral state classification

Videos were recorded using custom Matlab scripts at 10 Hz and compressed as .mp4 files using custom Matlab user interface (https://github.com/lachioma/FFMpeg-compress-videos) operating with the FFMpeg algorithm (Mathis and Warren 2018).

To visualize motion, we transformed the videos into motion-frames by computing the difference between consecutive frames. For creating the images in Fig. 5A, we generated an RGB image where the red and blue channels contained the values of frame n, while the green channel contained the values of frame n+1. This approach highlighted pixels with movement by showing them as pink or green (pink representing the pixels with the highest values in frame n, and green representing the corresponding pixels in frame n+1).

We manually categorized the motion frames into six different categories, which were resting, adjusting, grooming, walking, running, and sniffing+resting. For this purpose, we utilized a modified version of BENTO (https://github.com/karinmcode/bentoMAT), an open-source Matlab GUI for managing multimodal neuroscience data sets allowing browsing between recording sessions.

To streamline the labeling process, we employed a pre-labeling strategy that relied on the similarity between frames (https://github.com/karinmcode/bentoMAT/tree/master/bento/plugins/Prelabel_all_similar_frame s). Firstly, we computed the histogram of oriented gradients of motion frames (20 by 20 division of each frame) and stored the resulting values in a matrix of n frames by m HOG values (histogram of oriented gradients). Next, we reduced the size of the data by applying PCA (Principal Component Analysis) to the matrix. To obtain a 2D visualization (Fig. 5A), we used uMAP (Uniform Manifold Approximation and Projection) on the PCA outputs. Finally, we assigned a cluster ID to each frame using k-means clustering, with the option of using up to 12 clusters. All pre-labels were manually reviewed and reassigned to one of the six categories (Fig. 5A).

### Statistical analysis

Statistical analysis was performed using Matlab (Mathworks). Unless reported otherwise, all statistics were described as mean ± SEM. Statistical significance was defined as p<0.05 unless stated otherwise. The significance threshold was adjusted for multiple comparisons using the Bonferroni correction. One-sample Kolmogorov-Smirnov test served to determine the use of a non-parametric test. To investigate the proportions of neurons across all auditory cortical fields and pairwise comparisons, a Chi-square statistical test was used with a Bonferroni correction.

## Acknowledgements

We thank members of the Schneider lab for thoughtful feedback throughout this project. We extend our gratitude to Jessica Guevara for her expertise in animal care and technical assistance, and to Carina Sun for her assistance in labeling video frames. We thank Alessandro La Chioma for his assistance in setting up the software for the epifluorescence imaging. This research was supported by the Swiss National Science Foundation (K.M.); the National Institutes of Health (1R01-DC018802 to DMS); a Career Award at the Scientific Interface from the Burroughs Wellcome Fund (D.M.S); fellowships from the Searle Scholars Program, the Alfred P. Sloan Foundation, and the McKnight Foundation (D.M.S.); and an investigator award from the New York Stem Cell Foundation (D.M.S). D.M.S. is a New York Stem Cell Foundation - Robertson Neuroscience Investigator.

